# The loss of the PDIM/PGL virulence lipids causes differential secretion of ESX-1 substrates in *Mycobacterium marinum*

**DOI:** 10.1101/2024.01.09.574891

**Authors:** Bradley S. Jones, Daniel D. Hu, Kathleen R. Nicholson, Rachel M. Cronin, Matthew M Champion, Patricia A. Champion

## Abstract

The mycobacterial cell envelope is a major virulence determinant in pathogenic mycobacteria. Specific outer lipids play roles in pathogenesis, modulating the immune system and promoting the secretion of virulence factors. ESX-1 (ESAT-6 system-1) is a conserved protein secretion system required for mycobacterial pathogenesis (1, 2). Previous studies revealed that mycobacterial strains lacking the outer lipid PDIM have impaired ESX-1 function during laboratory growth and infection (3-5). The mechanisms underlying changes in ESX-1 function are unknown. We used a proteo-genetic approach to measure PDIM and PGL-dependent protein secretion in *M. marinum*, a non-tubercular mycobacterial pathogen that causes tuberculosis-like disease in ectothermic animals (6, 7). Importantly, *M. marinum* is a well-established model for mycobacterial pathogenesis (8, 9). Our findings showed that *M. marinum* strains without PDIM and PGL showed specific, significant reductions in protein secretion compared to the WT and complemented strains. We recently established a hierarchy for the secretion of ESX-1 substrates in four (I-IV) groups (10). Loss of PDIM differentially impacted secretion of Groups III and IV ESX-1 substrates, which are likely the effectors of pathogenesis. Our data suggests that the altered secretion of specific ESX-1 substrates is responsible for the observed ESX-1-related effects in PDIM-deficient strains.

**Importance:** *Mycobacterium tuberculosis*, the cause of human tuberculosis, killed an estimated 1.3 million people in 2022. Non-tubercular mycobacterial species are causing acute and chronic human infections. Understanding how these bacteria cause disease is critical. Lipids in the cell envelope are essential for mycobacteria to interact with the host and promote disease. Strains lacking outer lipids are attenuated for infection, but the reasons are unclear. Our research aims to identify a mechanism for attenuation of mycobacterial strains without the PDIM and PGL outer lipids in *M. marinum*. These findings will enhance our understanding of the importance of lipids in pathogenesis, and how these lipids contribute to other established virulence mechanisms.

## Introduction

Phthiocerol dimycocerosate (PDIM) and phenolic glycolipid (PGL) are outer lipid virulence factors in the mycolate outer membrane (MOM) (11-13). Although the precise role of PDIM/PGL in mycobacterial virulence remains elusive, PDIM/PGL support cell envelope structure, serve as a permeability barrier, enhance antibiotic tolerance, and directly modulate the immune system (14-18). These lipids recruit macrophages permissive for bacterial growth and survival while promoting evasion of bactericidal macrophages (11, 13, 17, 19-22).

PDIM and PGL are synthesized from cytoplasmic fatty acids via a series of discrete steps by the PpsA-E/FadD26, Mas/FadD28 and PapA5 enzymes (23-25). Their transport across the cytoplasmic membrane (CM) and mycobacterial arabinogalactan peptidoglycan complex (mAGP) relies on the MmpL7 and DrrABC transporters, and the LppX lipoprotein (11, 13, 26-29), although the precise mechanisms remain unknown.

Mycobacteria secrete proteins across the MOM to the cell surface and into the surrounding environment both during laboratory growth and infection (30-34). Proteins are exported across the CM by the Sec and SecA2 pathways, twin arginine transporter (TAT), and ESAT-6 (ESX) systems (35, 36). Yet, the mechanisms used by proteins to cross the impermeable mAGP and the MOM (37-39) are elusive. Recent studies suggest that PE/PPE and Esx (WXG_100) proteins facilitate protein translocation across the MOM (10, 40, 41).

Early during infection, mycobacterial secreted proteins promote interaction with host membranes and macrophage signaling pathways. The ESX-1 system secretes proteins that contribute to phagosomal damage (42-44), leading to pathogen release, along with its DNA, RNA and secreted proteins, into the macrophage cytoplasm (22, 44-46). Secreted proteins in the cytoplasm hinder phagosomal repair, affect host trafficking and promote dissemination of infected macrophages in the lung (44, 47-52). Mycobacterial DNA in the cytoplasm triggers the Type I IFN response (43, 45, 46, 53), promoting macrophage cell death and bacterial spread (43, 54).

In the absence of PDIM, ESX-1 function is reduced, affecting the Type I IFN response and phagosomal lysis (3, 4). It is unclear whether PDIM is required for ESX-1 protein secretion across the MOM (3) or for ESX-1 substrate function within the phagosome (4). Our study uses a proteogenetic approach (55) to quantify how PDIM and PGL influence protein secretion from *Mycobacterium marinum*. Our studies provide an explanation for the ESX-1 phenotypes in the absence of PDIM, suggesting a broader role of PDIM and PGL outer lipids in translocating mycobacterial proteins across the MOM.

## Results

### Generation and characterization of PDIM deficient *M. marinum* strains

The PDIM/PGL biosynthetic pathway is conserved in both *M. tuberculosis* and *M. marinum* (Figure 1A). We previously constructed *M. marinum* strains with disrupted PDIM production due to deletion of the *mas* gene (56) or the second ER domain of the *ppsC* gene (57). We generated an *M. marinum* strain with an unmarked deletion of the *drrABC* genes. PpsA-E/FadD26 synthesize phthiocerol; loss of the PpsCER domain ablates activity and abrogates PDIM production (19, 57-59). Mas/FadD28 synthesize mycocerosic acid; deletion of *mas* halts PDIM production (12, 23). The two PDIM components are joined by PapA5 and transported by DrrABC, MmpL7 and LppX (12, 25-27). Deletion of *drrABC* abolishes PDIM production and transport (12). Complementation strains were generated by introducing integrating plasmids with the *ppsC*_*ER*_ domain, the *drrABC* operon or the *mas* gene behind constitutive promoters into each deletion strain. We confirmed the resulting strains using PCR (Fig. S1) followed by targeted DNA sequencing, and qRT-PCR to measure transcription in the deletion and complementation strains (Fig. 1B and Fig. S2A and B). Deletion of the *mas* or *drrABC* genes abolished expression relative to the WT and complemented strains. Deletion of the *ppsC*_*ER*_ domain did not impact *ppsC* expression, as expected (57).

**Figure 1.**
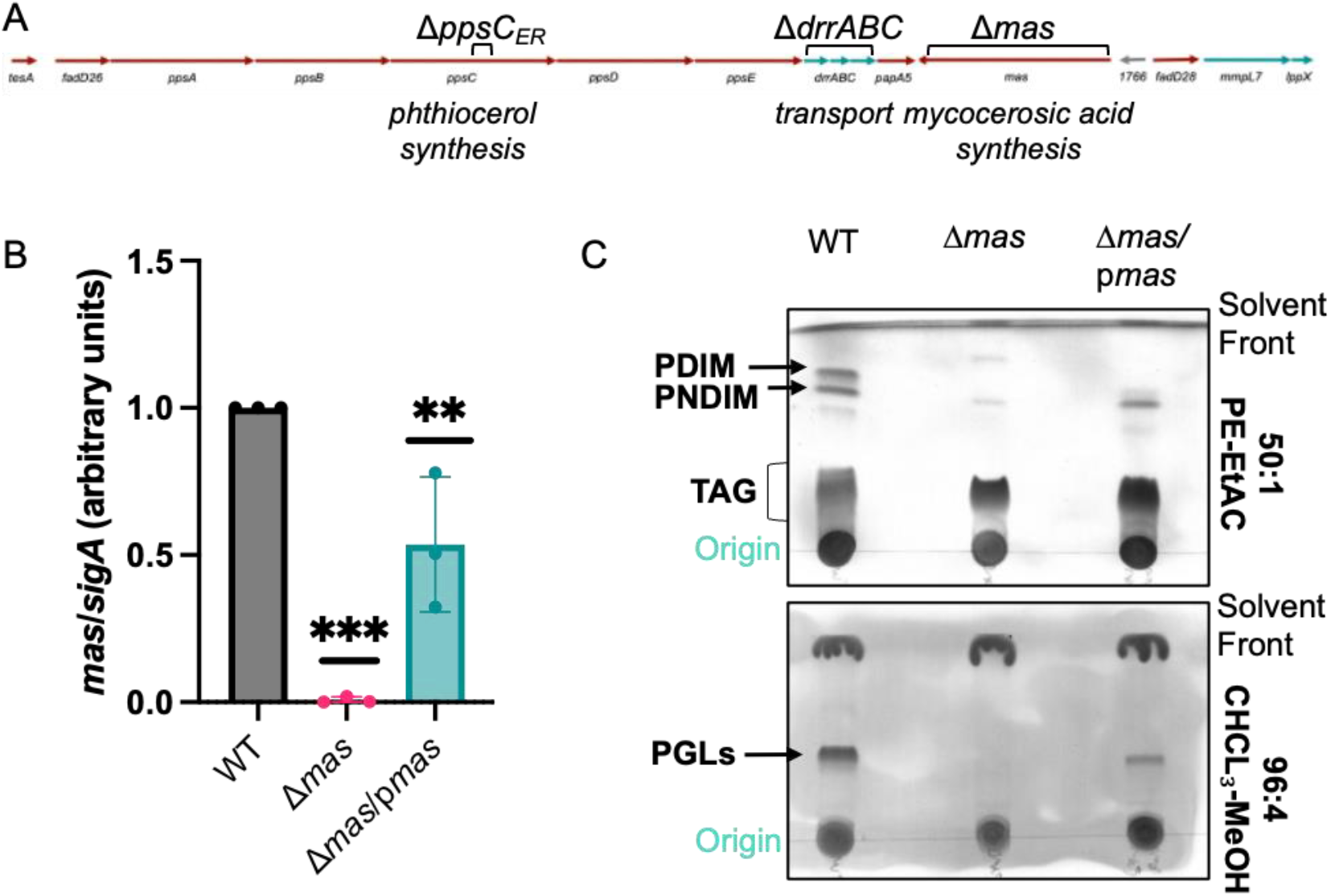
Characterization of PDIM/PGL deficient *M. marinum* strains. A. Schematic of the genetic loci required for PDIM production and transport. B. qRT-PCR of *mas* transcript relative to the *sigA* transcript compared to the WT *M. marinum* strain. Each datapoint is an independent biological replicate, and an average of three technical replicates. Statistical analysis was performed using an ordinary one-way ANOVA (*P*=.0003), followed by a Dunnett’s multiple comparison test, *** *P*=.0002, ** *P*=.0093. C. TLC of total lipids isolated from the WT, Δ*mas* and complemented *M. marinum* strains. 9 μl of total lipids was analyzed. This TLC is representative of at least three independent biological replicates.

We validated PDIM and PGL loss in each deletion strain (Fig. 1C and Fig. S2C) by isolating total soluble lipids and visualizing PDIM and PGL using thin layer chromatography. All three *M. marinum* deletion strains exhibited a lack of PDIM and PGL (Fig. 1C and Fig S2C). Reintroducing the *mas* gene restored PDIM in the Δ*mas* strain (Fig. 1C). PDIM and PGL were not complemented by restoring gene expression in the Δ*ppsC*_*ER*_ and Δ*drrABC* strains (Fig. S2C).

### The loss of PDIM/PGL has a general effect on mycobacterial protein secretion

Past studies linked ESX-1 secretion and PDIM/PGL, proposing that PDIM is necessary for ESX-1 protein secretion (3). To clarify PDIM/PGL’s role in mycobacterial protein secretion, we isolated secreted and cell associated protein fractions and quantified the protein levels in the mutant and complemented strains compared to the WT *M. marinum* strain using Label Free Quantitative (LFQ) Proteomics (Dataset S1). We measured ESX-1-dependent protein secretion as a control.

Deletion of the *eccCb*_*1*_ gene reduced ESX-1 substrate production and secretion (Fig. 2, top). We reasoned that if PDIM was specifically required for ESX-1 secretion, then deletion of the *mas, ppsC*_*ER*_ or *drrABC* genes will abolish ESX-1 secretion, similar to the Δ*eccCb*_*1*_ strain. Deletion of the *mas* gene (or deletion of the *drrABC* genes or the ER domain from the *ppsC* gene) resulted in a significant decrease in the levels of most secreted proteins (Fig. 2, middle and Fig. S3). Restoring *mas* expression in the Δ*mas* strain complemented the secretion defect (Fig. 2, bottom). These data support that PDIM is important for overall protein secretion from *M. marinum* during laboratory growth.

**Figure 2.**
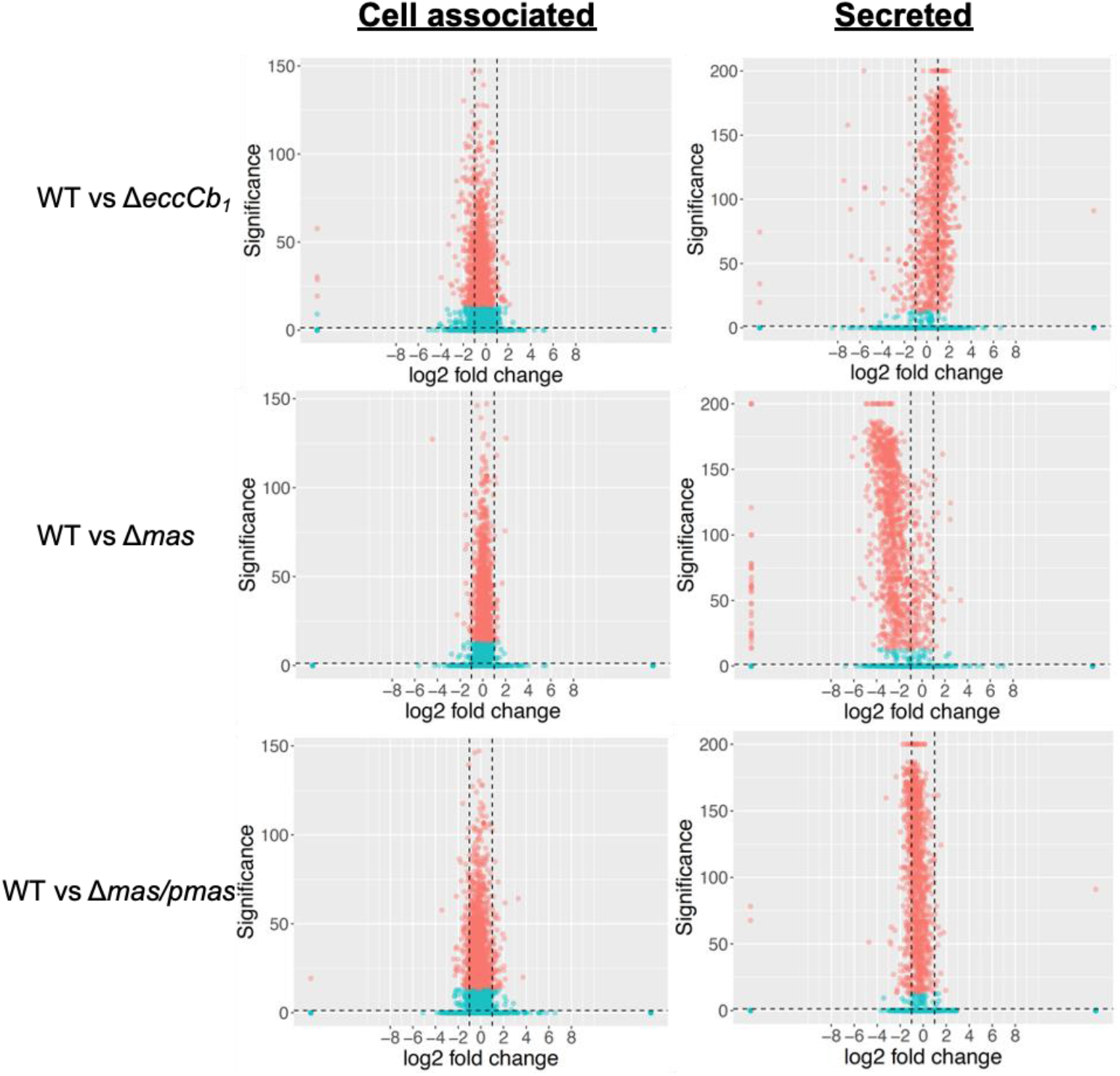
Loss of PDIM/ PGL results in widespread changes to protein secretion from *M. marinum*. Volcano plots of the secreted protein levels from cell associated (left) and secreted protein fractions from *M. marinum*. Values are the average of four biological replicates. Significance was plotted against the log2 fold change compared to the WT strain. blue dots indicate *P* ≤ 0.05; red dots indicate *P* > 0.05. Vertical, black dashed lines signify a log2 foldchange = +/- 1.

### The loss of PDIM/PGL specifically impacts the secretion of a subset of ESX-1 substrates

We previously demonstrated that ESX-1 substrates are hierarchically secreted from *M. marinum*. We categorized them into at least four groups of ESX-1 substrates (Fig. 3A, (10)). Group 1 includes EsxA, EsxB, PPE68 and MMAR_2894, which are essential for the secretion of the other ESX-1 substrates (10, 60, 61). Group 2 substrates include EspB, EspJ and EspK (10, 60, 62). These substrates are not required for the secretion of Group 1 substrates, but are essential for the secretion of ESX-1 Group 2 substrates (10). ESX-1 Group 3 substrates include EspE and EspF, and these substrates are dispensable for the secretion of the other ESX-1 substrate classes. Importantly, deletion of the Group 3 substrates is sufficient to abrogate the hemolytic activity of *M. marinum* (63).

**Figure 3.**
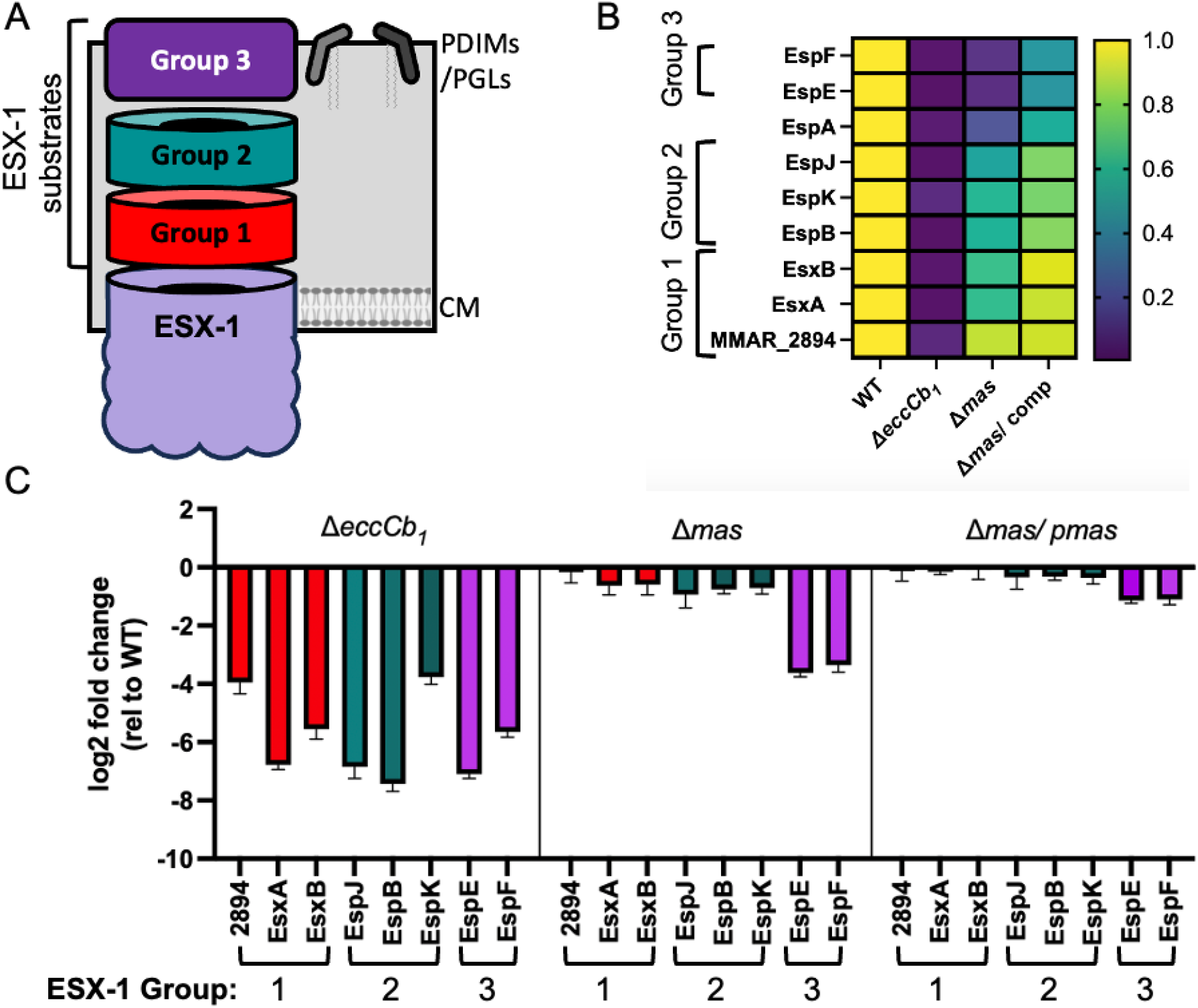
The loss of PDIM/PGL impacts ESX-1 secretion differentially. **A**. Schematic of the ESX-1 secretion hierarchy. ESX-1 represents the ESX-1 membrane components. Group 1: EsxB, EsxA, PPE68 and MMAR_2894, Group 2: EspB, EspK, EspJ, Group 3: EspE and EspF, from (10). **B**. Log2 Fold-Change of ESX-1 substrate levels in the secreted protein fractions compared to the WT strain. Data in Dataset 1C. **C**. Heat map of the ESX-1 substrate levels in the secreted protein fractions normalized to the WT strains (Data in Dataset 1D, from Dataset S1B)

Figure 3B and 3C indicates that the loss of PDIM specifically impacts the secretion of ESX-1 Group 3 substrates. In the Δ*eccCb*_*1*_ strain, ESX-1 substrate levels were significantly reduced (Figure S4, Dataset S1) due to feedback control of substrate gene transcription in the absence of the ESX-1 system (64). However, in the Δ*mas* strain, the substrate levels were not significantly reduced (Fig. S4), indicating no feedback control of substrate gene transcription due to the loss of the PDIM/PGL lipids. While ESX-1 substrate secretion was significantly reduced from the Δ*eccCb*_*1*_ strain because EccCb_1_ is directly required for substrate secretion (1, 65, 66), the levels of ESX-1 substrates secreted from the Δ*mas* strain differed, aligning with previously established substrate groups (Fig. 3B and C, Dataset S1). The Group 3 substrates (EspE, EspF) were the most affected by PDIM/PGL loss. The reduced secretion of Group 3 substrates were complemented by expression of *mas* in the Δ*mas* strain (Fig. 3B and 3C). Together, these data support that the loss of PDIM/ PGL differentially impacts the secretion of the known ESX-1 substrates.

### The absence of PDIM reduces ESX-1 function

Prior research demonstrated that PDIM-deficient *M. marinum* strain exhibited lower ESX-1-dependent hemolytic activity (5). EspE and EspF secretion were most impacted by PDIM/PGL loss. Deleting *espE* or *espF* from *M. marinum* abrogated hemolytic activity and attenuated *M. marinum* during macrophage infection (63). Because deleting *espE* or *espF* did not significantly affect the secretion of the additional known ESX-1 substrates, EspE and EspF are at the top of the hierarchy (10). In Figure 4A, WT *M. marinum* lysed sheep red blood cells (sRBCs) in a contact dependent, ESX-1 dependent manner. The Δ*eccCb*_*1*_ strain which lacks ESX-1 secretion, had significantly less hemolytic activity than the WT strain (*P<*.0001) (10, 61, 63). The Δ*mas* strain exhibited intermediate hemolytic activity, between the WT (*P*<.0001) and Δ*eccCb*_*1*_ strains (*P*<.0001), similar to a prior report (5). Reintroducing the *mas* gene in the Δ*mas* strain significantly increased hemolysis (*P*<.0001), but not to WT levels (WT vs Δ*mas*/p*mas, P*<.0001).

**Figure 4.**
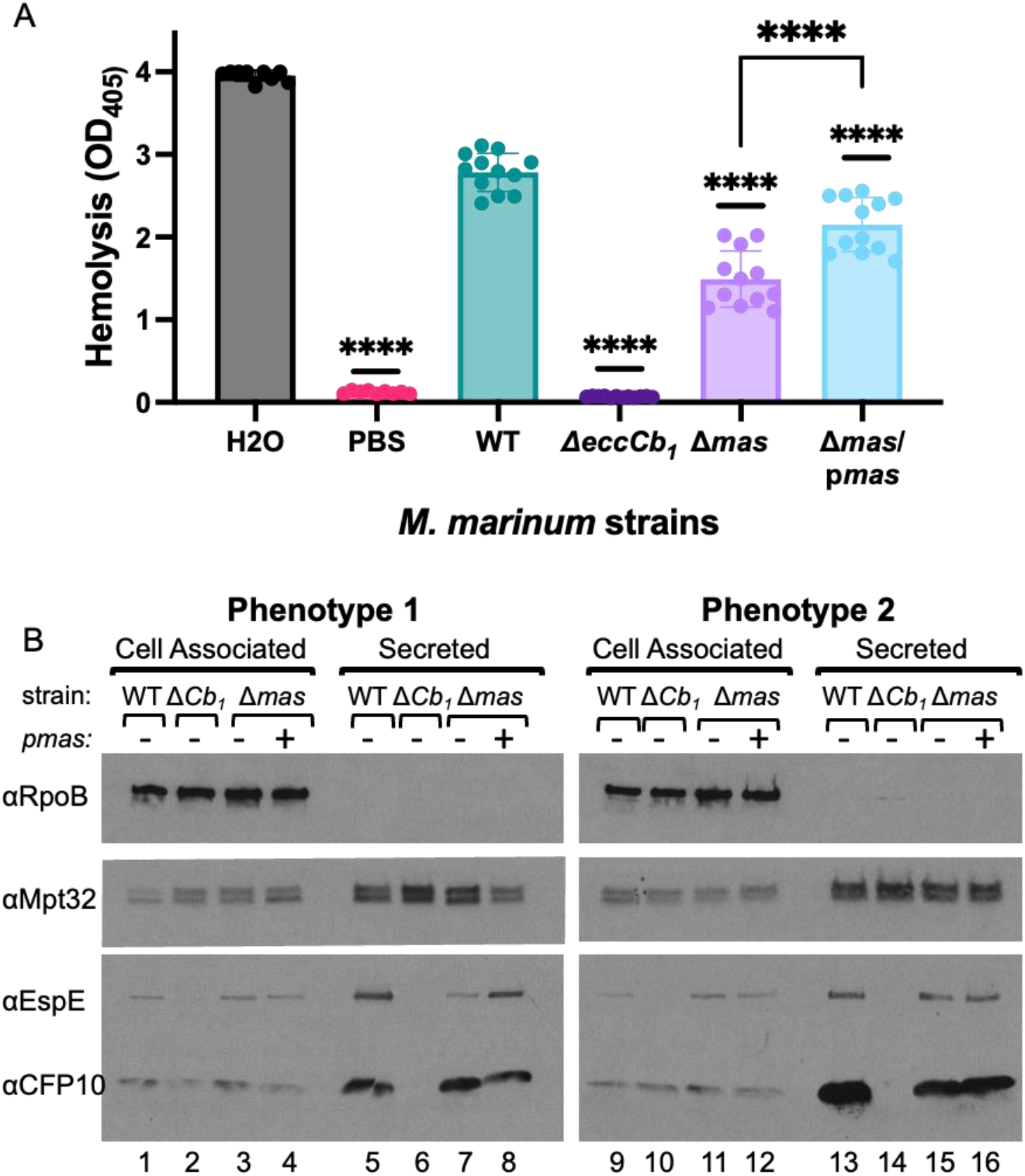
The loss of PDIM/PGL impacts hemolysis and the secretion of the EspE substrate. **A**. Hemolytic activity of *M. marinum*. Each data point is a single technical replicate, making up four biological replicates. Significance was determined using a one-way ANOVA, *P*<.0001, followed by a Tukey’s multiple comparison test **** *P*<.0001. **B**. Western blot analysis of *M. marinum* cell associated and secreted protein fractions. 10 μg of protein were loaded in each lane. RpoB is a control for lysis, MPT32, a protein secreted by the Sec system, is a loading control. The two blots shown were representative of at least four biological replicates.

We next used western blot analysis to examine ESX-1 substrate secretion. EsxB (CFP-10) was made (Fig. 4B, lane 1) and secreted (lane 4) from the WT strain. EsxB production was reduced (lane 2) and it was not secreted (lane 6) from the Δ*eccCb*_*1*_ strain, as previously reported (57). EsxB production (lanes 3 and 4) and secretion (lanes 7 and 8) were not noticeably impacted in the Δ*mas* and Δ*mas* complemented strains. EspE was produced (lane 1) and secreted (lane 5) from the WT strain. EspE levels were reduced in (lane 2) and was not secreted from the Δ*eccCb*_*1*_ strain (lane 6). Although EspE was made to WT levels in the Δ*mas* strain (lane 3), EspE secretion was reduced (lane 7). EspE secretion was restored in Δ*mas*/p*mas* strain (lane 8). These data are consistent with the LFQ proteomics data in Figures 2 and 3. Interestingly, as we replicated the western blot analysis, we sometimes observed no visible secretion phenotype, as shown in Figure 4, right. The LFQ analysis did not reveal WT levels of EspE and EspF in any of the replicates we tested. From these data, we conclude that western blot analysis, which is not quantitative, may lack the sensitivity to detect intermediate but biologically relevant changes in ESX-1 protein secretion, as supported by our prior work (10, 60).

## Discussion

This study aimed to understand the impact of the outer lipids PDIM and PGL on ESX-1 secretion. We measured a broad and significant reduction in *M. marinum* protein secretion in the absence of PDIM and PGLs. Interestingly, the loss of PDIM and PGLs differentially impacted ESX-1 protein secretion, particularly affecting the secretion of the Group 3 substrates, EspE and EspF. Our findings support that the ESX-1-associated phenotypes in strains lacking PDIM/ PGL are due to the reduced secretion of specific ESX-1 substrates.

At least three publications previously linked PDIM to ESX-1 secretion or function (3-5). Barczak et al. concluded PDIM is required for ESX-1 substrate secretion in *M. tuberculosis*. Deletion of *drrC* abrogated EsxA (ESAT-6) and EsxB (CFP-10) secretion, and the ESX-5 substrate, EspN, during laboratory growth. Other PDIM deficient strains in this study had WT-levels of EsxB secretion. The loss of ESX-1 secretion led to reduced IFN induction during macrophage infection with the Δ*drrC* strain as compared to the WT and complemented strains. IFN induction requires ESX-1 induced lysis of the phagosome membrane (44, 53). They concluded that the PDIM lipid at the cell surface was important for ESX-mediated protein secretion (3).

In Barczak et al. an intermediate secretion signal may have been missed because the amount of protein loaded in that study was based on the culture density as measured by OD_600_(3). We normalized the protein levels based on protein concentration as measured by BCA assay, because the OD_600_ of *M. marinum* grown Sauton’s media with low Tween-80 is an unreliable measure in our hands. Using this approach, we did not observe significant differences in the secretion of the Group 1 substrates, which includes EsxB, by western blot analysis or using LFQ proteomics. Importantly, Barczak et al. observed variability in EsxB secretion by different PDIM deficient strains (Δ*drrC* vs Δ*drrA*). Our study revealed that reduced EspE secretion was variable when detected by western blot analysis, but not by proteomics. We think the inconsistency in measuring these secretory phenotypes by western blot likely stems from their intermediate nature.

Quigley et al. connected PDIM transport to ESX-1 dependent events during macrophage infection, including phagosome lysis and the resulting macrophage cell death (4). They showed significantly reduced phagosomal lysis of an Δ*mmpL7 M. tuberculosis* strain compared to the WT and complemented strains. Aligned with our studies, Quigley et al. loaded 10μg of protein, found no defect in EsxA secretion by western blot analysis.

Osman et al. demonstrated that PDIM transport was important for phagosomal lysis, finding reduced phagosomal lysis of an Δ*mmpL7 M. marinum* strain during macrophage infection (5). Although the Δ*mmpL7* strain exhibited intermediate hemolytic activity between the ΔRD1 and WT strains, their data did not meet statistical significance. They suggested that PDIM enhanced the ESX-1 lytic activity, but offered that PDIM may contribute differently to hemolysis and phagosomal lysis. Our results show that *M. marinum* strains lacking PDIM/PGL production (Δ*mas* or Δ*ppsC*) or transport (Δ*drrABC*) have significantly reduced hemolysis compared the WT strain, and significantly higher hemolysis compared to the Δ*eccCb*_*1*_ *M. marinum* strain. We assessed hemolysis 90 minutes after initiating contact between *M. marinum* and the sRBCs, while Osman et al measured hemolysis after two hours (5). We previously found intermediate hemolysis phenotypes are most apparent at earlier timepoints (64, 67). We suspect that the intermediate hemolytic activity of the Δ*mmpL7* strain would have reached significance relative to WT strain at earlier timepoints.

An underlying commonality in all of these studies is that in strains lacking PDIM or PGL, ESX-1 activity is reduced, as reflected in hemolytic activity or phagosomal lysis during infection. Notably, at the time of these manuscripts, the secretory relationship between ESX-1 substrates was unclear. However, we now know that EspE/EspF secretion is essential for hemolysis, and for pathogenesis in macrophages (63). We have likewise defined a hierarchical secretory relationship between the ESX-1 substrates (10). Here we showed that the loss of PDIM specifically reduced EspE/EspF secretion from *M. marinum*. Therefore, we hypothesize that reduced EspE/EspF secretion caused the reduced hemolytic and phagolytic activity observed in the absence of PDIM. It is tempting to speculate that the secretion of proteins that colocalize with PDIM and PGL in the mycobacterial envelope is impacted by the loss of these lipids in *M. marinum*. This idea requires further exploration beyond these initial studies.

Our unbiased approach to measuring protein secretion revealed that losing PDIM/PGL broadly impacts protein secretion. Contrary to expectations based on PDIM’s role in maintaining envelope impermeability, making the cell envelope permeable did not increase protein secretion by increasing cellular lysis. Instead, overall protein secretion in the culture supernatants decreased, despite evidence of cellular lysis. Our study has direct implications for mycobacterial pathogenesis, as PDIM and PGL deficient *Mycobacterium* are attenuated in cellular infection models (11, 13, 58). Further study is needed to understand if the observed phenotypes due to the loss of mycobacterial outer lipids stem from changes in the secreted proteome.

## Methods

### Growth of Bacterial Strains

*Mycobacterium marinum* strains were derived from *Mycobacterium marinum* M strain (ATCC BAA-535). *M. marinum* strains were maintained at 30°C in either Middlebrook 7H9 Broth Base (Sigma Aldrich) supplemented with 0.5% glycerol and 0.1% Tween-80 (Fisher Scientific) or Middlebrook 7H11 Agar (Sigma Aldrich) plates supplemented with 0.5% glucose and 0.5% glycerol. Where appropriate, strains were supplemented with 50 μg/mL hygromycin B (Corning, Corning, NY), 20 μg/mL kanamycin (IBI Scientific, Dubuque, IA), and 60 μg/mL X-Gal (RPI, Mount Prospect, IL). *Escherichia coli* DH5α strains were grown in Luria-Bertani Broth (VWR) at 37°C and supplemented with 200 μg/mL hygromycin and 50 μg/mL kanamycin, where appropriate.

### Construction of *M. marinum* strains

The *ΔdrrABC* strain was generated using allelic exchange using the p2NIL vector [Addgene plasmid number 20188; a gift from Tanya Parish (68)] and the pGOAL19 vector [Addgene plasmid number 20190; a gift from Tanya Parish (68)], as previously published (61, 63, 64, 67, 69). Primers were purchased from Integrated DNA Technologies (IDT, Coralville, IA) and are listed in Table S2. The complementation plasmids were generated using FastCloning (70) using *M. marinum* M genomic DNA. Plasmids were introduced into *M. marinum* using electroporation as described previously (61, 63, 64, 67, 69, 71). Strains and plasmids are listed in Supplementary Tables S1. Plasmids and genetic deletions were confirmed by targeted DNA sequencing performed by the Notre Dame Genomics and Bioinformatics Facility.

### Thin-Layer Chromatography of PDIM and PGL Lipids

5 mL cultures of *M. marinum* in 7H9 defined broth (Millipore Sigma), and grown for 2-3 days until turbid, diluted into 50mL of 7H9 defined broth to an OD600 of 0.05 and grown for 72 hours. Mycobacterial cells were harvested via centrifugation (4000 rpm for 10 minutes). The resulting cell pellets were washed three times with phosphate-buffered saline (PBS). Lipids were extracted using a chloroform (Ricca Chemical Company, Arlington, TX) :methanol (Millipore Sigma) mixture (2:1) overnight in a fume hood. Overnight extraction mixtures were filtered through 6 mm Whatman filter paper (Millipore Sigma) using a porcelain Buchner funnel before separating aqueous and organic phase layers. Extracted lipids were moved to a clean glass vial and dried overnight in a fume hood under a gentle stream of air. Dried mycobacterial lipids were resuspended in a 100 μl mixture of chloroform: methanol (2:1) prior to analysis.

9 μl of total mycobacterial lipids were spotted onto aluminum-backed silica TLC plates (Millipore Sigma) and dried. To separate mycobacterial phthiocerol dimycocerosates (PDIM) and triacylglycerols (TAG), spotted lipids were migrated three times sequentially in a running solution of 50:1 petroleum ether (Sigma Aldrich) and ethyl acetate (Sigma Aldrich). The TLC plate was sprayed with phosphomolybdic acid (Millipore Sigma) and charred via heat gun (Westward) to visualize lipids.

Mycobacterial phenolic glycolipids (PGLs) were visualized using 9μl of total mycobacterial lipids migrated once in a solution of 96:4 chloroform and methanol. The TLC plate was sprayed with a solution of 1% α-naphthol (TCI Chemicals, Portland, OR) and charred via heat gun to visualize lipids.

### Hemolysis Assays

Hemolysis assays were performed exactly as described (10) except that sheep red blood cell and bacteria were incubated at 30**°**C for one and a half hours, rather than two hours.

### Secretion Assays

Cell-associated and secreted protein fractions were prepared from *M. marinum* exactly as previously described (10). Briefly, *M. marinum* bacterial cultures were grown in 7H9 defined media, diluted into 50 mL of Sauton’s media at an OD600 of 0.8 and grown for 48 hours before harvesting cellular and secreted fractions by centrifugation. Culture supernatants were filtered through 0.2 μm Nalgene Rapid-Flow Bottle Top Filters with PES membranes (Thermo Scientific) and concentrated using 15 mL, 3kDa Amicon Ultra Centrifugal Filter Units (Millipore Sigma). Mycobacterial cell pellets were resuspended in 0.5 mL of PBS, lysed via bead-beating with a BioSpec Mini-Beadbeater-24, and clarified by centrifugation. Protein concentrations of the resulting fractions were measured Pierce Micro BCA Protein Assay Kit (Thermo Scientific).

### Western Blot Analysis

10 μg of each protein sample was loaded into 4-20% TGX Gradient Gels (Bio-Rad) for analysis. Antibodies were diluted in 5% non-fat dry milk (RPI) in PBS with 0.1% Tween-20 (VWR). Rpoβ (anti-RNA polymerase beta mouse monoclonal antibody [clone: 8RB13]; VWR) was diluted 1:20,000. The following reagents were obtained through BEI resources, NIAID, NIH: polyclonal anti-*Mycobacterium tuberculosis* CFP-10 (gene Rv3874; antiserum, rabbit; NR-13801) and polyclonal anti-*Mycobacterium tuberculosis* Mpt-32 (gene Rv1860; antiserum, rabbit; NR-13807). CFP-10 was used at a 1:5,000 dilution. Mpt-32 was used at a 1:30,000 dilution. The EspE antibody (1:5,000 dilution) was a custom rabbit polyclonal antibody against the CGQQATLVSDKKEDD peptide (Genscript).

### Proteomics/LC-MS

LC-MS pure reagents (water, ethanol, acetonitrile, and methanol) were purchased from J.T. Baker (Radnor, PA). Iodoacetamide (IAA) was purchased from MP Biomedicals (Solon. OH). All other reagents are from Sigma-Aldrich (St. Louis, MO) unless specified. S-Trap mini devices were from Protifi (Huntington, NY). Trypsin Gold was purchased from Promega (Madison, WI). Hydrophilic–lipophilic balance (HLB) solid phase extraction (SPE) cartridges (1 cc/10 mg) from Waters (Milford, MA) were used to desalt peptide samples prior to analysis on a timsTOF Pro from Bruker Scientific LLC (Billerica, MA). Protein and peptide identification including label-free quantitation was performed using the PEAKS Online X search engine from Bioinformatics Solutions Inc (Waterloo, ON) (build 1.4.2020-10-21_171258).

Cell-associated and secreted cell lysate samples were prepared for LC-MS analysis as described (72, 73). 50 μg of each sample was prepared in 100 mM triethylammonium bicarbonate (TEAB), 5% sodium dodecyl sulfate (SDS), and 100 mM tris(2-carboxyethyl)phosphine (TCEP). Samples were heated for 10 minutes at 95°C, cooled, then IAA was added to 100 mM IAA for 30 minutes in dark.

Samples were acidified with *o-*phosphoric acid to a final concentration of 1.2% in 50 μL, then flocculated with 350 μL Binding Buffer containing 90% methanol and 10% 1M TEAB. Samples were passed through S-Trap Micro filters and followed by two 80 μL washes of Binding Buffer and 80 μL of 1:1 methanol/chloroform solution. A new collection tube was added, and 1 μg of trypsin in 80 μL was added. Samples were wrapped in Parafilm to prevent evaporative loss and incubated at 37°C for 12 hours. Digested peptides were spun through the filter, followed by two 80 μL elutions with 0.1% formic acid in water, and one 80 μL elution with 0.1% formic acid in 50% acetonitrile. Eluted peptides were vacuum concentrated for 20 minutes (to remove acetonitrile), and then desalted using 1cc/10 mg HLB SPE cartridges following manufacturer’s specifications, and dried by vacuum concentrator prior to analysis.

Desalted peptides were resuspended in 0.1% formic acid and water, to 1 mg/mL concentration. 100 ng of each sample was injected in triplicate onto an Evosep One and timsTOF Pro LC-MS system. Each sample was prepared in biological triplicate and technical triplicate. 15spd (Sample-per-day) methods were used on a 150 μm x 150 mm PepSep column with C_18_ ReproSil AQ stationary phase at 1.9mm particle size, 120Å pore size (Manufacturer’s protocol). Samples were loaded using Evotips following manufacturer’s specifications. Nano-ESI was used with a spray voltage of 1700V. MS was set to Parallel Accumulation, Serial Fragmentation Data Dependent Mode (PASEF-DDA) with a mass range of 100–1700 m/z, ion mobility range of 0.6–1.6 v*s/cm^2^, and ramp and accumulation times of 100 ms. Each precursor consisted of 10 PASEF ramps for a cycle time of 1.17 s. Precursors were filtered to contain only charges from 2 to 5. MS/MS collision energy settings were set to ramp from 20 eV at 0.6 ion mobility to 70 eV at 1.6 ion mobility. Instrument tune parameters were set to default for proteomic studies with the following differences: quadrupole low mass set to 20 m/z, focus pre-TOF pre-pulse storage set to 5 ms.

Protein and peptide identification and label-free quantitation were performed using the PEAKS Online X search engine. Search database used was the *M. marinum* proteome from Mycobrowser (v4 release). Search settings were set to manufacturer defaults unless specified. 3 missed cleavages were allowed, at a semi-specific search. Fixed modification included carbamidomethylation of cysteine, while variable modifications included protein N-terminal acetylation, deamidation of asparagine and glutamine, pyroglutamic acid from glutamine and glutamic acid, and oxidation of methionine. All peptide-spectrum matches were filtered to a 1% false discovery rate. Cell-associated samples were normalized by total ion current (TIC). RAW files are deposited and available at MassIVE/Proteome Exchange ftp://MSV000093713@massive.ucsd.edu (password PDIM2024#) MSV000093713.

LFQ search results were trimmed as follows. Technical replicate values for each sample were averaged if at least two of the three replicates were nonzero values. The same averaging logic was applied to biological replicates. Only nonzero values were averaged. All mutant samples were ratioed the corresponding WT values, to generate fold-change ratios.

### RNA extraction

*M. marinum* strains were grown as described in the Secretion Assays methods section. 15 mL of Sauton’s culture were pelleted by centrifugation. Bacterial pellets were resuspended in Qiagen RLT buffer (Qiagen, Hilden, Germany) supplemented with 1% β-mercaptoethanol. Lysates were generated by bead beating with the Biospec Mini-BeadBeater-24 and clarified with centrifugation. Total RNA was extracted using the RNeasy Mini Kit (Qiagen), according to manufacturer’s instructions.

### qRT-PCR

RNA (500 ng) was treated with RQ1 DNase (Promega) according to the manufacturer’s instructions. 1 μl of DNase treated RNA was converted to cDNA using random hexamers (IDT) and SuperScript II Reverse Transcriptase (Invitrogen) according to manufacturer instructions. cDNA was quantified using a Nanodrop 2000 (Thermo Scientific, Waltham, MA). 10 μl qRT-PCR reactions were prepped with 250 ng of cDNA, SYBR™ Green PCR Master Mix (Applied Biosystems, Carlsbad, CA) and 1 μM of each oligonucleotide. Reactions were run using Applied Biosystems MicroAmp Fast 96-well plates (0.1 ml) (Life Technologies). Plates were run on QuantStudio 3 Real-Time PCR system (Thermo Fisher). Cycle conditions were as follows: 50 °C for 2 min., 95 °C for 10 min. then 40 cycles at 95 °C for 15 sec and 60 °C for 1 min. Then, a dissociation step of 95 °C for 15 sec., 60 °C for 1 min., 95 °C for 15 sec., and 60 °C for 15 sec.

qRT-PCR reactions were analyzed using ΔΔCt comparisons. qRT-PCR results were normalized to WT transcript abundance using the following equations:

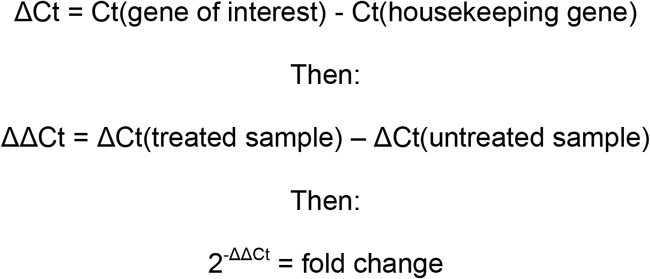

### R Code for Volcano Plots

>WT_vs_dmas$threshold=as.factor(WT_vs_dmas$Significance < 13) *Determine which Proteins were significantly changed, 13 is the 95% percentile or 10*^*-1*.*3*^

>g <-ggplot(data = WT_vs_dmas, aes(x=L2FC, y = Significance, color = threshold)) + geom_point(alpha = 0.4, size = 1.75) + xlim(c(−8, 8)) + xlab(“log2 fold change”) + (ylab(“-log10 p-value”)+ theme_bw() + theme(legend.position = “none”))

>g

>g +theme(text=element_text(size = 20))+ geom_hline(yintercept = -log10(0.05), linetype = “dashed”) + geom_vline(xintercept = c(log2(0.5), log2(2)), linetype = “dashed”)+scale_x_continuous(breaks = c(seq(−8,8,2)))

## Acknowledgments

K.R.N and B.J. were supported by an Eck Institute for Global Health Fellowship. K.R.N. was also supported by an Arthur J. Schmitt Fellowship. P.A.C. is supported by the National Institutes of Health under award numbers AI156229, AI106872, AI149147, and AI149235. M.M.C. is supported by the National Institutes of Health under award numbers GM144372 and GM139277. We thank the Mass Spectrometry and Proteomics Facility at Notre Dame. The content of this article is solely our responsibility and does not necessarily represent the official views of the National Institutes of Health.

## References

1. S. A. Stanley, S. Raghavan, W. W. Hwang, J. S. Cox, Acute infection and macrophage subversion by Mycobacterium tuberculosis require a specialized secretion system. Proc Natl Acad Sci U S A 100, 13001–13006 (2003).

2. T. Hsu et al., The primary mechanism of attenuation of bacillus Calmette-Guerin is a loss of secreted lytic function required for invasion of lung interstitial tissue. Proc Natl Acad Sci U S A 100, 12420–12425 (2003).

3. A. K. Barczak et al., Systematic, multiparametric analysis of Mycobacterium tuberculosis intracellular infection offers insight into coordinated virulence. PLoS Pathog 13, e1006363 (2017).

4. J. Quigley et al., The Cell Wall Lipid PDIM Contributes to Phagosomal Escape and Host Cell Exit of Mycobacterium tuberculosis. MBio 8 (2017).

5. M. M. Osman, A. J. Pagan, J. K. Shanahan, L. Ramakrishnan, Mycobacterium marinum phthiocerol dimycocerosates enhance macrophage phagosomal permeabilization and membrane damage. PLoS One 15, e0233252 (2020).

6. F. Linell, A. Norden, Mycobacterium balnei, a new acid-fast bacillus occurring in swimming pools and capable of producing skin lesions in humans. Acta Tuberc Scand Suppl 33, 1–84 (1954).

7. H. F. Clark, C. C. Shepard, Effect of Environmental Temperatures on Infection with Mycobacterium Marinum (Balnei) of Mice and a Number of Poikilothermic Species. J Bacteriol 86, 1057–1069 (1963).

8. T. C. Pozos, L. Ramakrishnan, New models for the study of Mycobacterium-host interactions. Curr Opin Immunol 16, 499–505 (2004).

9. A. E. Chirakos, A. Balaram, W. Conrad, P. A. Champion, Modeling Tubercular ESX-1 Secretion Using Mycobacterium marinum. Microbiol Mol Biol Rev 84 (2020).

10. R. M. Cronin, M. J. Ferrell, C. W. Cahir, M. M. Champion, P. A. Champion, Proteo-genetic analysis reveals clear hierarchy of ESX-1 secretion in Mycobacterium marinum. Proc Natl Acad Sci U S A 119, e2123100119 (2022).

11. L. R. Camacho, D. Ensergueix, E. Perez, B. Gicquel, C. Guilhot, Identification of a virulence gene cluster of Mycobacterium tuberculosis by signature-tagged transposon mutagenesis. Mol Microbiol 34, 257–267 (1999).

12. L. R. Camacho et al., Analysis of the phthiocerol dimycocerosate locus of Mycobacterium tuberculosis. Evidence that this lipid is involved in the cell wall permeability barrier. J Biol Chem 276, 19845–19854 (2001).

13. J. S. Cox, B. Chen, M. McNeil, W. R. Jacobs, Jr., Complex lipid determines tissuespecific replication of Mycobacterium tuberculosis in mice. Nature 402, 79–83 (1999).

14. A. M. Block et al., Mycobacterium tuberculosis Requires the Outer Membrane Lipid Phthiocerol Dimycocerosate for Starvation-Induced Antibiotic Tolerance. mSystems 8, e0069922 (2023).

15. M. S. Siegrist, C. R. Bertozzi, Mycobacterial lipid logic. Cell Host Microbe 15, 1–2 (2014).

16. S. A. Stanley, J. S. Cox, Host-Pathogen Interactions During Mycobacterium tuberculosis infections. Curr Top Microbiol Immunol 374, 211–241 (2013).

17. C. J. Cambier et al., Mycobacteria manipulate macrophage recruitment through coordinated use of membrane lipids. Nature 505, 218–222 (2014).

18. C. Passemar et al., Multiple deletions in the polyketide synthase gene repertoire of Mycobacterium tuberculosis reveal functional overlap of cell envelope lipids in hostpathogen interactions. Cell Microbiol 16, 195–213 (2014).

19. C. J. Cambier, S. M. O’Leary, M. P. O’Sullivan, J. Keane, L. Ramakrishnan, Phenolic Glycolipid Facilitates Mycobacterial Escape from Microbicidal Tissue-Resident Macrophages. Immunity 47, 552–565 e554 (2017).

20. M. A. Kirksey et al., Spontaneous phthiocerol dimycocerosate-deficient variants of Mycobacterium tuberculosis are susceptible to gamma interferon-mediated immunity. Infect Immun 79, 2829–2838 (2011).

21. T. A. Day et al., Mycobacterium tuberculosis strains lacking surface lipid phthiocerol dimycocerosate are susceptible to killing by an early innate host response. Infect Immun 82, 5214–5222 (2014).

22. Y. Cheng, J. S. Schorey, Mycobacterium tuberculosis-induced IFN-beta production requires cytosolic DNA and RNA sensing pathways. J Exp Med 215, 2919–2935 (2018).

23. A. K. Azad, T. D. Sirakova, L. M. Rogers, P. E. Kolattukudy, Targeted replacement of the mycocerosic acid synthase gene in Mycobacterium bovis BCG produces a mutant that lacks mycosides. Proc Natl Acad Sci U S A 93, 4787–4792 (1996).

24. A. K. Azad, T. D. Sirakova, N. D. Fernandes, P. E. Kolattukudy, Gene knockout reveals a novel gene cluster for the synthesis of a class of cell wall lipids unique to pathogenic mycobacteria. J Biol Chem 272, 16741–16745 (1997).

25. O. A. Trivedi et al., Dissecting the mechanism and assembly of a complex virulence mycobacterial lipid. Mol Cell 17, 631–643 (2005).

26. M. Jain, J. S. Cox, Interaction between polyketide synthase and transporter suggests coupled synthesis and export of virulence lipid in M. tuberculosis. PLoS Pathog 1, e2 (2005).

27. G. Sulzenbacher et al., LppX is a lipoprotein required for the translocation of phthiocerol dimycocerosates to the surface of Mycobacterium tuberculosis. EMBO J 25, 1436–1444 (2006).

28. M. H. Touchette, J. C. Seeliger, Transport of outer membrane lipids in mycobacteria. Biochim Biophys Acta 10.1016/j.bbalip.2017.01.005 (2017).

29. M. H. Touchette et al., A Screen for Protein-Protein Interactions in Live Mycobacteria Reveals a Functional Link between the Virulence-Associated Lipid Transporter LprG and the Mycolyltransferase Antigen 85A. ACS Infect Dis 3, 336–348 (2017).

30. C. Abou-Zeid et al., The secreted antigens of Mycobacterium tuberculosis and their relationship to those recognized by the available antibodies. J Gen Microbiol 134, 531–538 (1988).

31. M. G. Sonnenberg, J. T. Belisle, Definition of Mycobacterium tuberculosis culture filtrate proteins by two-dimensional polyacrylamide gel electrophoresis, N-terminal amino acid sequencing, and electrospray mass spectrometry. Infect Immun 65, 4515–4524 (1997).

32. Y. Ge et al., Top down characterization of secreted proteins from Mycobacterium tuberculosis by electron capture dissociation mass spectrometry. J Am Soc Mass Spectrom 14, 253–261 (2003).

33. E. F. Perkowski et al., The EXIT Strategy: an Approach for Identifying Bacterial Proteins Exported during Host Infection. MBio 8 (2017).

34. F. Sayes et al., Multiplexed Quantitation of Intraphagocyte Mycobacterium tuberculosis Secreted Protein Effectors. Cell Rep 23, 1072–1084 (2018).

35. L. S. Ligon, J. D. Hayden, M. Braunstein, The ins and outs of Mycobacterium tuberculosis protein export. Tuberculosis (Edinb) 92, 121–132 (2012).

36. C. Chagnot, M. A. Zorgani, T. Astruc, M. Desvaux, Proteinaceous determinants of surface colonization in bacteria: bacterial adhesion and biofilm formation from a protein secretion perspective. Front Microbiol 4, 303 (2013).

37. P. A. Lambert, Cellular impermeability and uptake of biocides and antibiotics in Grampositive bacteria and mycobacteria. J Appl Microbiol 92 Suppl, 46S–54S (2002).

38. A. H. Delcour, Outer membrane permeability and antibiotic resistance. Biochim Biophys Acta 1794, 808–816 (2009).

39. P. S. Manzanillo et al., The ubiquitin ligase parkin mediates resistance to intracellular pathogens. Nature 501, 512–516 (2013).

40. L. S. Ates et al., Essential Role of the ESX-5 Secretion System in Outer Membrane Permeability of Pathogenic Mycobacteria. PLoS Genet 11, e1005190 (2015).

41. Q. Wang et al., PE/PPE proteins mediate nutrient transport across the outer membrane of Mycobacterium tuberculosis. Science 367, 1147–1151 (2020).

42. N. van der Wel et al., M. tuberculosis and M. leprae translocate from the phagolysosome to the cytosol in myeloid cells. Cell 129, 1287–1298 (2007).

43. R. Simeone et al., Phagosomal rupture by Mycobacterium tuberculosis results in toxicity and host cell death. PLoS Pathog 8, e1002507 (2012).

44. D. Houben et al., ESX-1-mediated translocation to the cytosol controls virulence of mycobacteria. Cell Microbiol 14, 1287–1298 (2012).

45. P. S. Manzanillo, M. U. Shiloh, D. A. Portnoy, J. S. Cox, Mycobacterium Tuberculosis Activates the DNA-Dependent Cytosolic Surveillance Pathway within Macrophages. Cell Host Microbe 11, 469–480 (2012).

46. R. O. Watson, P. S. Manzanillo, J. S. Cox, Extracellular M. tuberculosis DNA Targets Bacteria for Autophagy by Activating the Host DNA-Sensing Pathway. Cell 150, 803–815 (2012).

47. A. Coros, B. Callahan, E. Battaglioli, K. M. Derbyshire, The specialized secretory apparatus ESX-1 is essential for DNA transfer in Mycobacterium smegmatis. Mol Microbiol 69, 794–808 (2008).

48. M. S. Siegrist et al., Mycobacterial Esx-3 is required for mycobactin-mediated iron acquisition. Proc Natl Acad Sci U S A 106, 18792–18797 (2009).

49. S. R. Elliott, A. D. Tischler, Phosphate responsive regulation provides insights for ESX-5 function in Mycobacterium tuberculosis. Curr Genet 62, 759–763 (2016).

50. T. A. Gray et al., Intercellular communication and conjugation are mediated by ESX secretion systems in mycobacteria. Science 354, 347–350 (2016).

51. C. Portal-Celhay et al., Mycobacterium tuberculosis EsxH inhibits ESCRT-dependent CD4(+) T-cell activation. Nat Microbiol 2, 16232 (2016).

52. E. Mittal et al., Mycobacterium tuberculosis Type VII Secretion System Effectors Differentially Impact the ESCRT Endomembrane Damage Response. MBio 9, e01765–01718 (2018).

53. S. A. Stanley, J. E. Johndrow, P. Manzanillo, J. S. Cox, The Type I IFN response to infection with Mycobacterium tuberculosis requires ESX-1-mediated secretion and contributes to pathogenesis. J Immunol 178, 3143–3152 (2007).

54. D. G. Russell, The ins and outs of the Mycobacterium tuberculosis-containing vacuole. Cell Microbiol 18, 1065–1069 (2016).

55. K. R. Nicholson, C. B. Mousseau, M. M. Champion, P. A. Champion, The genetic proteome: Using genetics to inform the proteome of mycobacterial pathogens. PLoS Pathog 17, e1009124 (2021).

56. O. A. Collars et al., An N-acetyltransferase required for ESAT-6 N-terminal acetylation and virulence in Mycobacterium marinum. mBio 14, e0098723 (2023).

57. E. A. Williams et al., A Nonsense Mutation in Mycobacterium marinum That Is Suppressible by a Novel Mechanism. Infect Immun 85 (2017).

58. J. Yu et al., Both phthiocerol dimycocerosates and phenolic glycolipids are required for virulence of Mycobacterium marinum. Infect Immun 80, 1381–1389 (2012).

59. O. Vergnolle et al., Biosynthesis of cell envelope-associated phenolic glycolipids in Mycobacterium marinum. J Bacteriol 197, 1040–1050 (2015).

60. M. M. Champion, E. A. Williams, R. S. Pinapati, P. A. Champion, Correlation of Phenotypic Profiles Using Targeted Proteomics Identifies Mycobacterial Esx-1 Substrates. J Proteome Res 3, 5151–5164 (2014).

61. R. E. Bosserman, K. R. Nicholson, M. M. Champion, P. A. Champion, A New ESX-1 Substrate in Mycobacterium marinum That Is Required for Hemolysis but Not Host Cell Lysis. J Bacteriol 201 (2019).

62. B. McLaughlin et al., A mycobacterium ESX-1-secreted virulence factor with unique requirements for export. PLoS Pathog 3, e105 (2007).

63. A. E. Chirakos, K. R. Nicholson, A. Huffman, P. A. Champion, Conserved ESX-1 substrates EspE and EspF are virulence factors that regulate gene expression. Infect Immun 10.1128/IAI.00289-20 (2020).

64. R. E. Bosserman et al., WhiB6 regulation of ESX-1 gene expression is controlled by a negative feedback loop in Mycobacterium marinum. Proc Natl Acad Sci U S A 10.1073/pnas.1710167114 (2017).

65. K. M. Guinn et al., Individual RD1-region genes are required for export of ESAT-6/CFP-10 and for virulence of Mycobacterium tuberculosis. Mol Microbiol 51, 359–370 (2004).

66. L. Y. Gao et al., A mycobacterial virulence gene cluster extending RD1 is required for cytolysis, bacterial spreading and ESAT-6 secretion. Mol Microbiol 53, 1677–1693 (2004).

67. K. G. Sanchez et al., EspM Is a Conserved Transcription Factor That Regulates Gene Expression in Response to the ESX-1 System. mBio 11 (2020).

68. T. Parish, N. G. Stoker, Use of a flexible cassette method to generate a double unmarked Mycobacterium tuberculosis tlyA plcABC mutant by gene replacement. Microbiology 146 (Pt 8), 1969–1975 (2000).

69. R. E. Bosserman, C. R. Thompson, K. R. Nicholson, P. A. Champion, Esx Paralogs Are Functionally Equivalent to ESX-1 Proteins but Are Dispensable for Virulence in Mycobacterium marinum. J Bacteriol 200, e00726–00717 (2018).

70. C. Li et al., FastCloning: a highly simplified, purification-free, sequence- and ligationindependent PCR cloning method. BMC Biotechnol 11, 92 (2011).

71. T. Parish, Electroporation of Mycobacteria. Methods Mol Biol 2314, 273–284 (2021).

72. J. W. Saelens et al., An ancestral mycobacterial effector promotes dissemination of infection. Cell 10.1016/j.cell.2022.10.019 (2022).

73. O. A. Collars et al., An N-acetyltransferase required for EsxA N-terminal protein acetylation and virulence in Mycobacterium marinum. bioRxiv 10.1101/2023.03.14.532585, 2023.2003.2014.532585 (2023).

